# Longitudinal Resting-State fMRI of Awake Mice During Habituation: Stress, Head Motion, and Functional Connectivity

**DOI:** 10.64898/2025.12.22.696087

**Authors:** Sang-Han Choi, Geun Ho Im, Seong-Gi Kim

## Abstract

Awake mouse fMRI is a powerful tool for both neuroscience and translational research. To minimize head motion during scanning, habituation under physical restraint is commonly used. However, it remains unclear how stress levels and head motion evolve during habituation, particularly within the MRI environment. To address this, we repeatedly measured plasma corticosterone (CORT) levels in three groups of mice - controls, mice habituated outside the MRI magnet, and mice habituated within the fMRI environment - and acquired longitudinal resting-state fMRI data daily during an eight-day habituation period and again 15 days post-habituation at 15.2 T. We found that CORT levels initially increased by approximately twofold and gradually decreased during habituation outside the magnet, whereas in mice habituated within the fMRI environment, CORT levels increased two- to fourfold and remained elevated throughout the habituation period. One week after habituation, CORT levels returned to baseline in both groups. Throughout all resting-state fMRI scanning sessions, head motion and functional connectivity remained stable, likely due to the well-designed restraint cradle that permitted paw movement. These results suggest that additional habituation days do not further reduce stress, provided that head motion remains within acceptable limits.

## 1 Introduction

Mouse fMRI has become a valuable tool in neuroscience, owing to the vast availability of transgenic mouse models that enable precise investigations linking brain structure, function, and gene expression. It provides a platform for mapping cell-type-specific functional circuits across the whole brain and for bridging mechanistic insights from sophisticated mouse studies to human research through a shared MRI framework. Traditionally, anesthesia has been employed to minimize head motion and reduce stress during imaging[1]; however, it profoundly alters physiological and neural activity[2-14]. Because mice are particularly sensitive to anesthetic agents, maintaining stable anesthetic depth and acquiring reproducible fMRI data remain challenging[15]. As a result, awake fMRI approaches are increasingly being adopted to achieve more physiologically relevant measurements of brain function[16-22].

In awake mouse fMRI, minimizing head motion and stress is critical, as the two factors are likely interrelated. Head motion is typically reduced by restraining the head and body, which, in turn, induces stress. To mitigate both motion and stress, acclimation procedures are required. Although various acclimation approaches have been reported[20, 23-25], the best strategy would be to train awake mice within the actual MRI environment[18]. However, this approach is often impractical and unsustainable due to limited scanner availability and high operational costs.

As an alternative, habituation training is typically conducted in a mock scanner, followed by fMRI sessions in the real MRI environment. This approach presumes that the reduced stress achieved during habituation persists during actual scanning, although this is unlikely given the distinct sensory and vibrational characteristics of the MRI scanning environment. In a recent awake mouse fMRI study[25], plasma corticosterone levels (CORT)-an indicator of stress-returned close to the baseline level after two weeks of habituation (Fig. 2, RS2 & RS3). However, CORT increased to about 2-3 times the habituated value after the first MRI session and decreased slightly during the follow-up scanning session (Supp. 6 in Xu et al., 2022), suggesting that habituation inside the MRI environment itself may be necessary. However, it has not been systematically investigated whether habituation in an MRI environment effectively reduces stress and how this relates to head motion and brain functional connectivity.

To address these questions, we repeatedly measured plasma CORT levels in the same mice that were trained either outside or inside the MRI magnet and acquired longitudinal resting-state fMRI (rsfMRI) data from awake mice during an eight-day habituation period at 15.2 T. Because our newly developed cradle effectively minimizes head motion even without prior habituation[16], it was possible to perform repeated rsfMRI measurements daily both before habituation and after 1 to 7 days of acclimation. We found that plasma CORT levels initially increased by approximately 2-4 times relative to baseline following restraint, and gradually decreased during habituation outside the magnet, but not inside the MRI scanner. Head motion and functional connectivity remained consistent throughout the 8-day habituation period, suggesting that additional habituation does not further reduce stress as long as head motion remains within acceptable limits.

## 2 Materials and Methods

### 2.1 Animals

Thirty male C57BL/6 mice (10-12 weeks old) were used with approval from the Institutional Animal Care and Use Committee of Sungkyunkwan University. Because CORT levels are known to differ by sex[26-28], only male mice were included in this study. All experiments were performed in accordance with the standards for humane animal care and in compliance with the guidelines of the Animal Welfare Act and the National Institutes of Health Guide for the Care and Use of Laboratory Animals.

### 2.2 Head-post implantation

The head-post (Fig. 1A), made of acrylonitrile butadiene styrene (ABS), was 0.75-1.3 mm thick, and was slightly curved to ensure tight contact with the skull[16]. A detailed head-post implantation protocol is described in our previous publications[16, 29]. In short, animals were initially anesthetized with isoflurane (ISO) anesthesia at 3∼4%, then maintained at 1.5∼2.0% during the stereotaxic surgery for head-post placement. After placing the mouse in a stereotactic frame, the hair was shaved, and 2% lidocaine hydrochloride was subcutaneously injected under the scalp. All soft tissues were removed and cleaned with saline-soaked cotton buds. After ensuring the skull was completely clean, the ∼0.2-g head-post was placed on the exposed skull. All the exposed regions, including the head-post, were covered with a smooth layer of dental cement (Model #: C&B, Sun Medical Co, Shiga, Japan) 3∼4 times to reduce distortion in the MRI images caused by air. After the surgery, the mice were allowed to recover from anesthesia and then returned to their cages.

**Figure 1:**
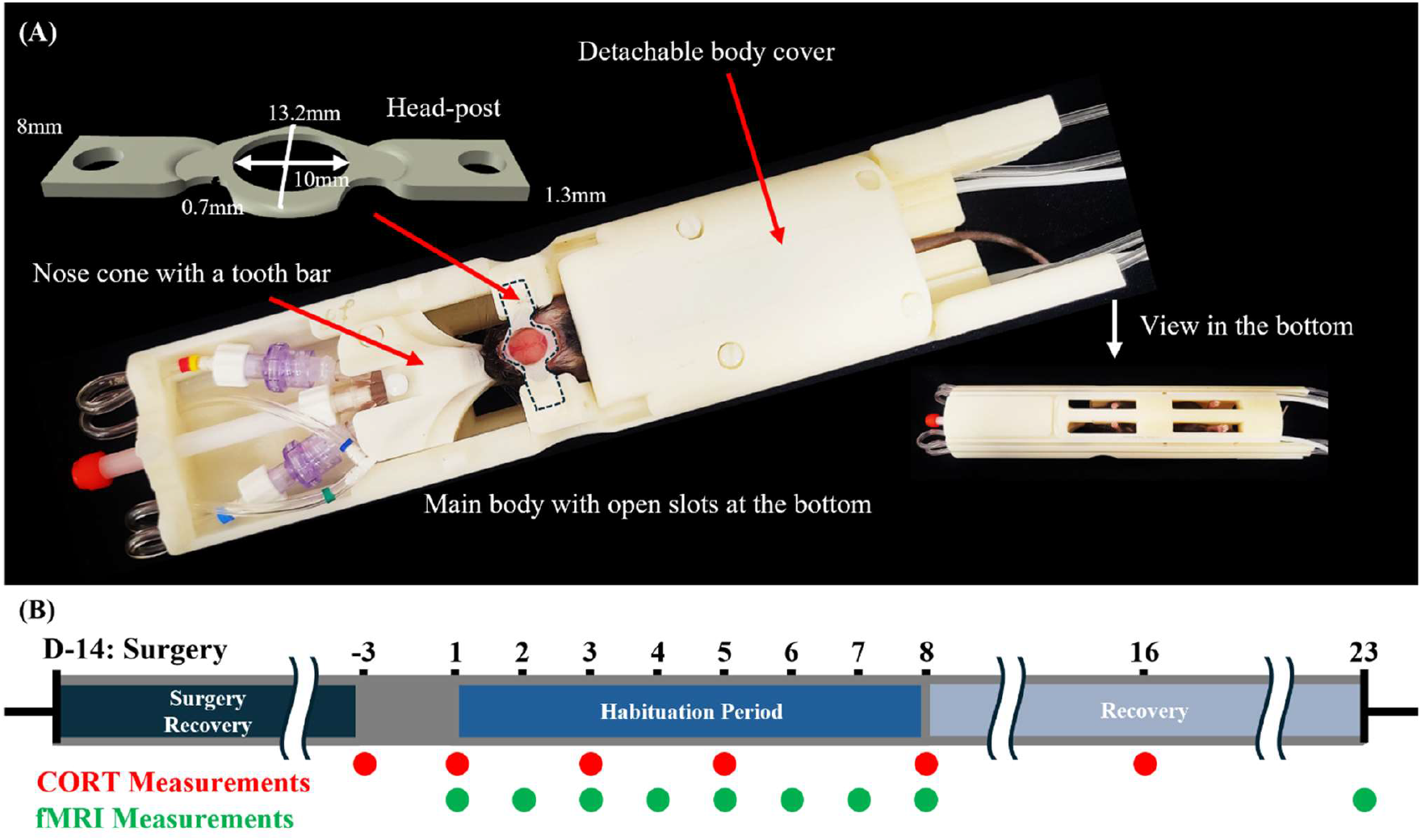
Experimental setup and protocol. **(A)** Cradle setup with a head-post design. **(B)** Experimental schedule for habitation and CORT measurements. Three groups were used: control, cradle habituation outside the magnet, and the fMRI group.

### 2.3 Overall habituation protocol

Two weeks after head-post implantation, mice were randomly assigned to three groups (n = 10 per group): no habituation (Group 1), habituation outside the magnet (Group 2), and habituation inside the magnet during fMRI scanning (Group 3). To monitor stress-related physiological changes, body weight was measured on days 1-8, 10 (2 days post-habituation), 12 (4 days post-habituation), 16 (8 days post-habituation), and 23 (15 days post-habituation), with day 1 corresponding to the first day of habituation.

To restrain each mouse in Groups 2 and 3, we used a custom-designed cradle with a body frame and a detachable body cover that conformed to the mouse’s body and included open slots at the bottom to allow paw movement (Fig. 1A), thereby reducing stress[16]. Isoflurane anesthesia was briefly administered to minimize stress during the positioning process, with induction at 5% ISO and maintenance at 2% ISO via a nose cone. After the mouse was positioned in the cradle, its mouth was secured to a tooth bar integrated into the nose cone. The entire procedure lasted less than 5 minutes. Daily 2-hour habituation or scanning sessions were then conducted under awake conditions on days 1-8, followed by an additional scanning session on day 23 (15 days post-habituation) (Fig. 1B).

For Group 2, MRI acoustic noise was not presented, as previous studies have shown that EPI scanning noise is not a major stressor in C57BL/6 mice[25, 30]. This allowed multiple mice to be habituated simultaneously without the need for a dedicated soundproof mock scanner room.

### 2.4 Plasma corticosterone concentration measurement

To repeatedly measure plasma CORT levels in the same mice, we followed the blood collection and CORT assay procedures described by Xu et al[25]. Plasma CORT concentrations were measured on days -3, 1, 3, 5, and 8 within 20 minutes after each habituation or scanning session for Groups 2 and 3, and again on day 16 (8 days post-habituation) to confirm the return of CORT levels to baseline (Fig. 1B). For Group 1 mice, CORT measurements were performed on the same schedule.

Briefly, under 1-1.5% isoflurane anesthesia, approximately 40 μL of blood was collected from the saphenous vein using heparin-coated glass capillaries sealed at one end (Paul Marienfeld GmbH & Co. KG, Lauda-Königshofen, Germany). One mouse in Group 3 died during blood sampling; thus, nine mice were included in the final analysis for that group.

Blood-filled capillaries were placed in 1.5 mL Eppendorf tubes on ice and centrifuged (Model: 5424R, Eppendorf, Hamburg, Germany) at 10,000 rpm for 10 minutes. The clear plasma layer (15-20 μL) was carefully transferred to PCR tubes (PCR-02-C, AxyGen, Union City, CA, USA) and stored at -20 °C until further analysis. After the completion of all experiments, plasma samples were analyzed for CORT concentration using an enzyme-linked immunosorbent assay (ELISA) kit (DRG-4164, DRG Instruments GmbH, Marburg, Germany).

### 2.5 MRI acquisition of awake mice at 15.2T (Group 3)

Two-hour MRI experiments were conducted daily on days 1-8 using a 15.2 T MRI system (BioSpec, Bruker, Ettlingen, Germany), with an additional fMRI session performed on day 23. The system was equipped with an 11 cm horizontal-bore magnet and an actively shielded gradient coil (6 cm inner diameter, maximum strength 100 G/cm, rise time 110 μs). A 12 mm inner-diameter surface coil was used for both transmission and reception. The mouse brain was positioned at the magnet isocenter, and field inhomogeneity was minimized using the MAPSHIM protocol with an ellipsoidal shim volume covering the cerebrum (ParaVision 360, Bruker BioSpin, USA).

Detailed MRI procedures have been described previously[16]. Anatomical images were acquired using a fast low-angle shot (FLASH) sequence with the following parameters: matrix size = 256 × 128, field of view (FOV) = 15.0 × 7.5 mm^2^ (0.059 × 0.059 mm^2^ in-plane resolution), 20 slices (0.5 mm thickness), repetition time/echo time (TR/TE) = 390/3.4 ms, and number of averages = 2.

Resting-state functional images were obtained using gradient-echo single-shot echo-planar imaging (EPI) with the following parameters: matrix size = 96 × 48, FOV = 15.0 × 7.5 mm^2^ (0.156 × 0.156 mm^2^ in-plane resolution), 20 slices (0.5 mm thickness), TR/TE = 1000/11.7 ms, and 600 volumes (10 min per run). Each daily session consisted of five 10-minute runs.

During all experiments, physiological signals were continuously monitored using a physiological monitoring system (Model 1030, SA Instruments, Inc., Stony Brook, NY, USA) and recorded using a data acquisition system (Biopac Systems, Inc., Goleta, CA, USA). Body motion and respiration rate were monitored using a pressure sensor (Model RS-301) placed on the ventral surface. Body temperature was monitored via a rectal probe and maintained at 37 °C using a circulating warm-water pad.

### 2.6 Respiration rate analysis from physiological data obtained during fMRI

Since respiration rate is indicative of stress level, respiration (breaths per minute, BPM) was estimated from pressure sensor signals recorded using the Biopac system, which captured respiration-induced chest movements. The average BPM for each animal was calculated from a 2000-s segment (approximately 33 min) acquired near the midpoint of each session by counting the sinusoidal respiratory cycles per minute in the signal.

### 2.7 Motion analysis from resting-state fMRI data

Slice-time-corrected rsfMRI data were used to estimate six head-motion parameters (three translations and three rotations) during the image realignment step. Subsequently, frame-wise displacement (FD) was calculated as a head-motion index by summing the absolute displacements of the six motion parameters, assuming a spherical mouse brain with a radius of 5 mm[31]. Among the 9 animals across 9 daily sessions, three sessions showed unusual motion along the phase-encoding direction, likely due to technical issues related to phase locking and unstable shimming in the Bruker software; therefore, these sessions were excluded from further analyses. For each run, an FD time course was generated. To assess the impact of habituation, motion histograms and mean FD values across days and runs were computed.

### 2.8 Resting-state fMRI data analyses

All FLASH anatomy data and BOLD time series EPI data for each mouse were analyzed using several tools and software packages, including the Analysis of Functional Neuroimages package[32], FMRIB Software Library (FSL), Advanced Normalization Tools (ANTs)[33], and Matlab codes (MathWorks, USA). EPI images were co-registered to anatomical images and subsequently to the Allen Mouse Brain Atlas, following procedures described in detail in our previous studies[16]. Briefly, individual EPI images were first aligned to their corresponding T_2_^*^-weighted anatomical images using linear transformation. These co-registered images were then normalized to a mouse brain template using nonlinear transformations and subsequently aligned to the Allen Mouse Brain Atlas.

Preprocessing of the acquired data included slice-timing correction, spike removal, motion correction, regression of motion components (12 regressors) as well as global and CSF signals, spatial smoothing with a 0.5 mm full width at half maximum, detrending, and band-pass filtering (0.01-0.25 Hz). The use of 12 motion parameters (six first-order parameters and their six temporal derivatives) followed prior studies on motion correction[18, 34]. CSF regions were automatically selected based on the criterion that voxel intensities exceeded half of the maximum intensity.

Although white matter signals are often regressed in rsfMRI analyses, we did not include them here because white matter could not be reliably identified in our data.

Brain region-specific ROIs for quantitative analysis were defined according to the Allen Mouse Brain Atlas (https://atlas.brain-map.org/). Three analyses were performed:

1. Seed-based correlation maps were generated to verify that the rsfMRI connectivity patterns were consistent with commonly reported networks in the literature. Seed ROIs were selected in the left primary somatosensory cortex (SSp) for the sensory network, the bilateral anterior cingulate cortex (ACC) for the default mode network (DMN), the left hippocampal region for the hippocampal network, and the left striatum for the caudate-putamen network.
2. Apparent functional connectivity (FC) specificity was evaluated by calculating the ratio of signal correlation between homologous SSp and the correlation between the SSp and the ACC[35].
3. FC analysis was conducted using twenty ROIs, with ten ROIs selected in each hemisphere: MOp (primary motor), SSp, VIS (visual), ACC, PL (prelimbic), ORB (orbitofrontal), RSP (retrosplenial), HIP (hippocampal), DORsm (thalamic sensory motor), and DORpm (thalamic polymodal). Correlation matrices were compared both across nine-day sessions and across repeated trials within each day.

### 2.9 Statistics

All results are presented as the mean standard deviation (SD). The statistical significance was tested using ANOVA and t tests. For all statistical tests, significance was considered according to an FDR-corrected p-value <0.05.

## 3 Results

### 3.1 Physiological parameters

To assess stress levels during the habituation process, we measured body weight and plasma CORT in three experimental groups, and respiratory rate in the fMRI group. Mean baseline body weights on day 1 were 27.1 ± 2.35 g (n = 10), 28.4 ± 2.36 g (n = 10), and 27.9 ± 3.35 g (n = 9) for the control (Group 1), outside-the-magnet habituation (Group 2), and fMRI habituation groups (Group 3), respectively (Fig. 2A). In the control group, body weight remained stable for the first 6 days and then gradually increased, although an earlier onset of weight gain had been expected. This delayed increase was likely due to stress associated with repeated blood sampling for CORT analysis. Both habituation groups showed a similar ∼10% loss of body weight during the 8-day habituation period, followed by weight gain after habituation ended, indicating comparable stress levels between the two groups.

**Figure 2:**
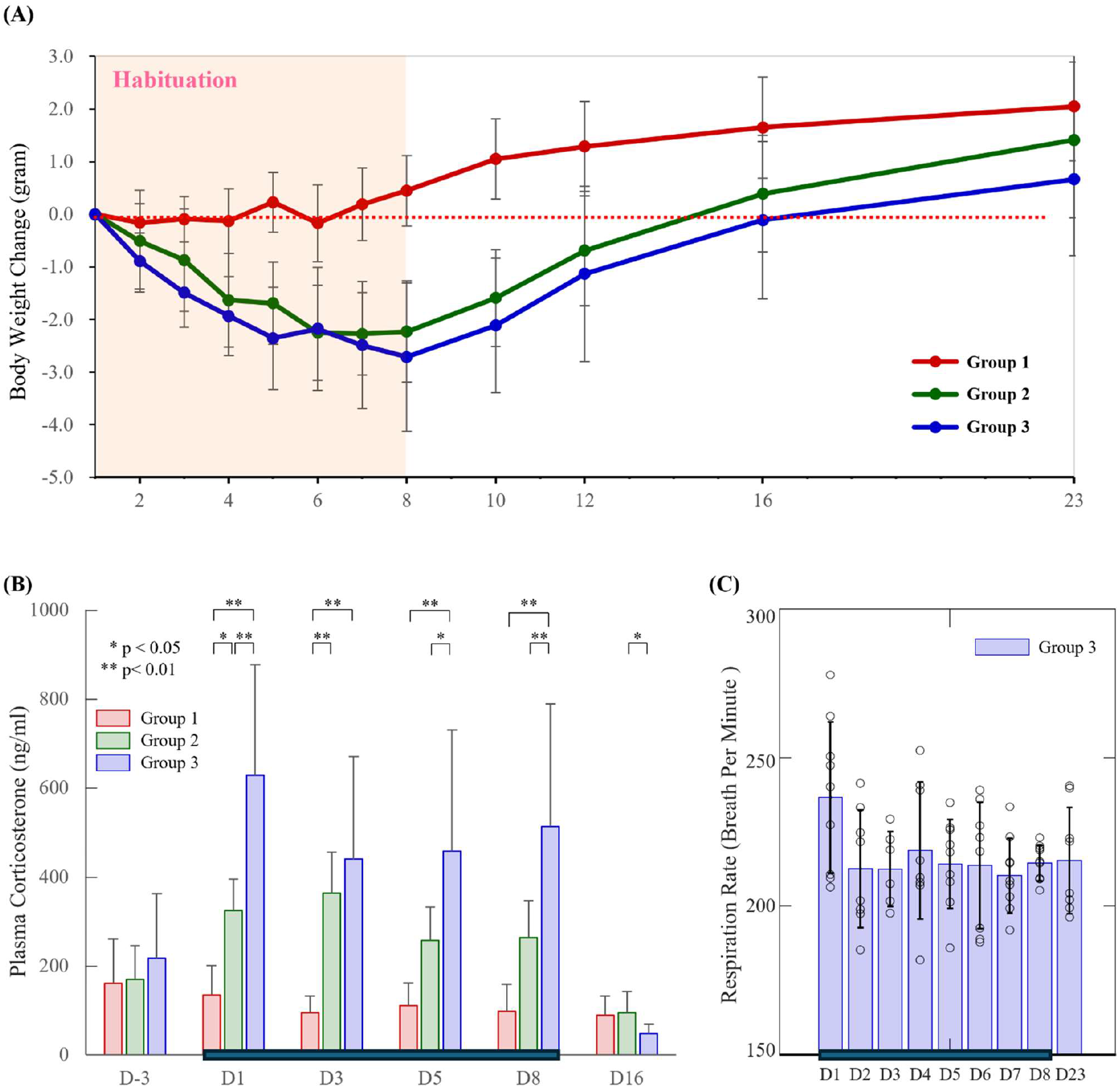
Stress level indices before, during and after the habituation period. Changes in body weight **(A)** and plasma corticosterone concentrations **(B)** under the three experimental conditions. Group 1: control without any habituation; Group 2: 2-hr restraint outside the magnet; and Group 3: 2-hr fMRI inside the 15.2T magnet. Statistical significance was assessed using ANOVA followed by post hoc tests. **(C)** Changes in breaths per minute for Group 3 during the fMRI scan period.Habituation was performed daily from day 1 to day 8 and is indicated by the horizontal bar. Day 16 corresponds to 8 days post-habituation, and day 23 corresponds to 15 days post-habituation.

CORT levels were measured before, during, and one week after the 8-day habituation period (Fig. 2B). Baseline CORT levels were 160.1 ± 100.5 ng/ml, 169.1 ± 75.7 ng/ml, and 216.9 ± 145.6 ng/ml (mean ± SD) for Group 1, Group 2, and Group 3, respectively. Two animals exhibited baseline CORT levels >400 ng/ml, underscoring the high sensitivity of CORT to handling stress. Habituation increased CORT levels by approximately twofold in the restraint group and two- to fourfold in the fMRI habituation group. The average CORT levels from days 1 - 8 were 108.8 ± 55.6 ng/ml (n = 10 animals × 4 days), 301.8 ± 89.3 ng/ml (n = 10 animals × 4 days), and 509.8 ± 257.0 ng/ml (n = 9 animals × 4 days) for Group 1, Group 2, and Group 3, respectively. During the habituation period, the fMRI habituation group consistently exhibited higher CORT levels than the restraint group, except on day 3. The highest CORT levels were observed on day 1 in the fMRI group. One week after the habituation procedures, CORT levels returned to baseline in all groups, indicating that the 8-day habituation regimen did not induce lasting chronic stress.

Respiratory rates in the magnet-habituation group were 238.6 ± 25.4 breaths per minute (BPM) during the first fMRI scan and averaged 216.5 ± 7.9 BPM across sessions (n = 74 days sessions in 9 animals). No significant differences were observed across days (Fig. 2C). Overall, respiratory rates remained stable at approximately 210-220 BPM, which falls within the normal range reported in the literature (210-270 BPM) [36-40]. This range is consistent with our previous observation using the same cradle setup (227.3 ± 28.3 BPM)[16], but lower than values obtained with an earlier version of the cradle (260-300 BPM)[18]. Despite exhibiting normal respiratory rates, awake mice undergoing fMRI scanning still exhibited signs of stress based on body weight loss and elevated CORT levels.

### 3.2 Head motion

Because stressed mice tend to move more, habituation is commonly used to reduce head motion, particularly in fMRI studies. In our study, although CORT levels remained elevated throughout the 8-day fMRI scanning period, it is still possible that habituation reduced motion during later sessions. Therefore, we compared FD across days and across runs within each session.

Figure 3A,C shows FD time courses averaged across five runs from nine animals on the first fMRI day (without prior habituation) and on day 8 (after seven habituation sessions). FD fluctuations were observed to a similar extent on both days. Histograms illustrate the distributions of FD values for all runs (600 volumes/run × 5 runs × 9 animals) on day 1 (Fig. 3B) and day 8 (Fig. 3D). The average FD values were 28.0 ± 21.1 μm and 33.8 ± 24.9 μm for day 1 and day 8, respectively, indicating that head motion during fMRI was comparable without or with one week of habituation.

**Figure 3:**
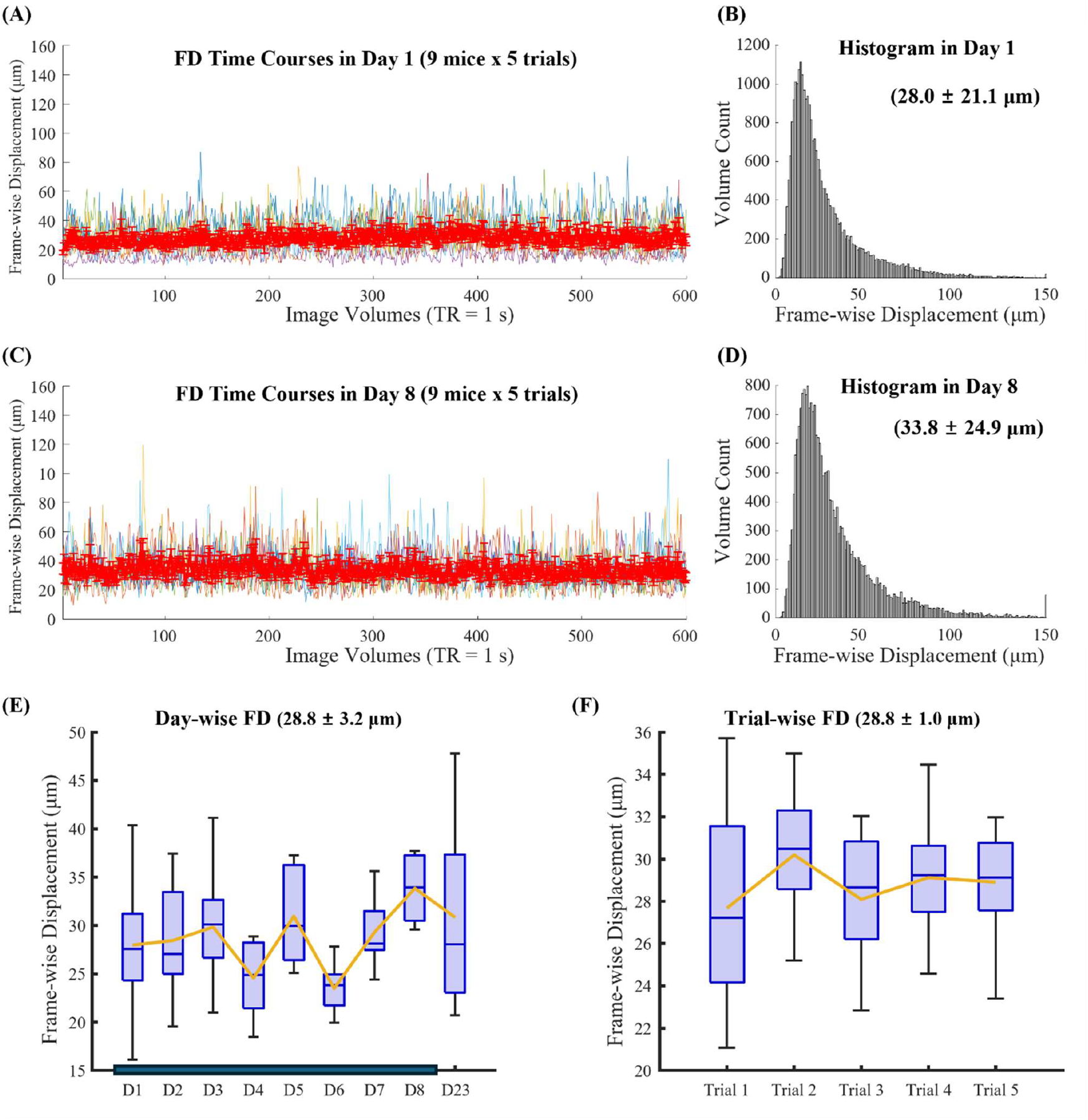
Frame-wise displacement (FD) representing head motion during resting-state fMRI in awake mice. Averaged FD time courses **(A**,**C)** and FD histogram **(B**,**D)** across five runs from nine animals are shown day 1 **(A**,**B)** and day 8 **(C**,**D)**. Day 1 corresponds to the first fMRI session without prior habituation, whereas day 8 corresponds to the final habituation session following seven days of habituation. Box plots summarize FD across scanning days **(E)** and across consecutive runs within each session **(F)**. Each session consisted of a 2-hour scan performed at 15.2 T.

Average FD values were plotted across days 1-8 and day 23 (15 days after the end of habituation) (Fig. 3E), as well as across the five consecutive runs within each session (Fig. 3F). The average FD value in day 8 was significantly higher than the average FD value in Day 4 (p-value = 0.02) and 6 (p-value = 0.01). No decreasing trends in head motion were observed across successive sessions or runs, indicating that head motion remained stable when the restraint system was well designed to reduce stress levels, such as by allowing free paw movements.

### 3.3 Resting-state fMRI

To assess whether rsfMRI maps were reliably detected in our data, correlation maps were generated from rsfMRI data on day 1 using seed regions in the bilateral ACC, left SSp, left hippocampal region, and left striatum. Correlation maps obtained from the ACC seed resembled the DMN, while maps generated from the SSp, hippocampal region, and striatum exhibited robust bilateral connectivity (see Fig. 4A,B for DMN and somatosensory network examples; and Suppl Fig. 1 and 2 for hippocampal region and striatum). These rsfMRI connectivity patterns closely resembled those reported in the literature[17].

**Figure 4:**
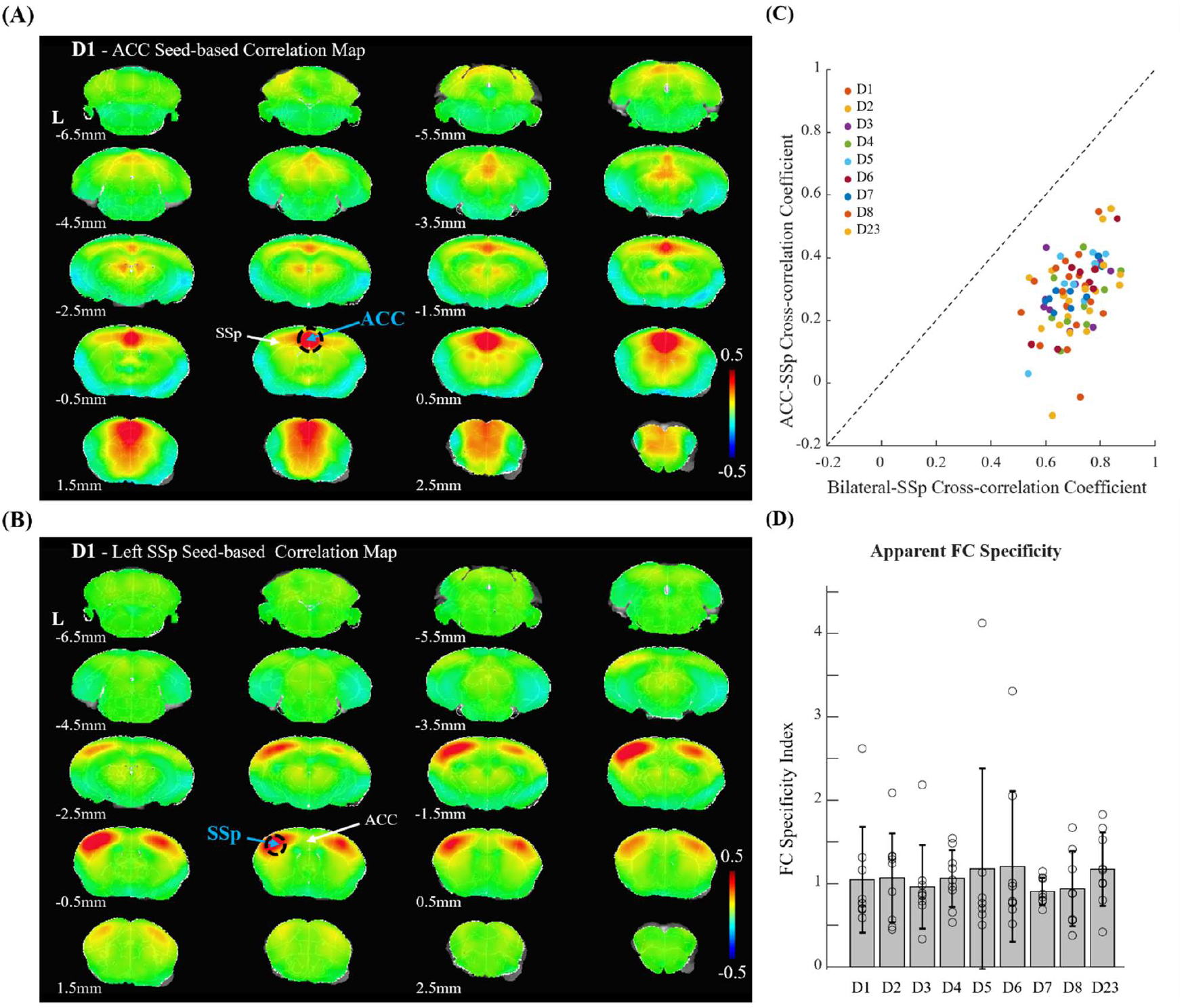
Seed-based cross-correlation maps and functional connectivity specificity index. **(A-B)** ACC- and SSp-seed based cross-correlation maps of the resting-state fMRI on day 1 showing DMN and somatosensory network, respectively. Similar connectivity patterns were observed in rs-fMRI datasets from other days. **(C)** Scatter plot of bilateral somatosensory (SSp-SSp) cross-correlation values versus ACC-SSp cross-correlation values for each animal and day session; a unity slope (slope = 1.0) line is included for reference. **(D)** Apparent function connectivity specificity across sessions. The specificity was calculated as the logarithm of the absolute value of [(L-SSp vs. R-SSp) × 2] / [(L-SSp vs. ACC) + (R-SSp vs. ACC)] where pairwise temporal cross-correlations between the left and right primary somatosensory (SSp) and anterior cingulate (ACC) areas were used.

To assess the specificity of FC, we adopted an index proposed by Grandjean et al[35], defined as the ratio of bilateral SSp connectivity to the correlation between the SSp and ACC. As expected, correlations between interhemispheric SSp were higher than those between SSp and the ACC (Fig. 4C; individual sessions). FC specificity was calculated the logarithm of the absolute value of the ratio between bilateral SSp and SSp-ACC correlation coefficients, and remained consistent across sessions with different levels of habituation (Fig. 4D). An FC specificity value of 1.0 indicates that bilateral SSp connectivity is approximately 2.7 times stronger than SSp-ACC connectivity.

We next examined whether resting-state functional connectivity changed during the habituation process, despite the similar CORT levels, head motion and FC specificity values observed across days. Cross-correlation analysis was performed among 20 ROIs in both hemispheres, with 10 ROIs per each hemisphere. Figure 5A shows the FC matrices from days 1 and 8. No obvious changes were detected over time (Fig. 5B). For improved visualization, −log(p-value) was plotted, where a p-value of 0.05 without FDR correction corresponds to ∼3.0. Inter-hemispheric functional connectivity, a hallmark feature of rsfMRI FC, was also evaluated across days (Fig. 5C), and no significant changes were observed, except the visual system. These results indicate that resting-state FC remained stable throughout habituation.

**Figure 5:**
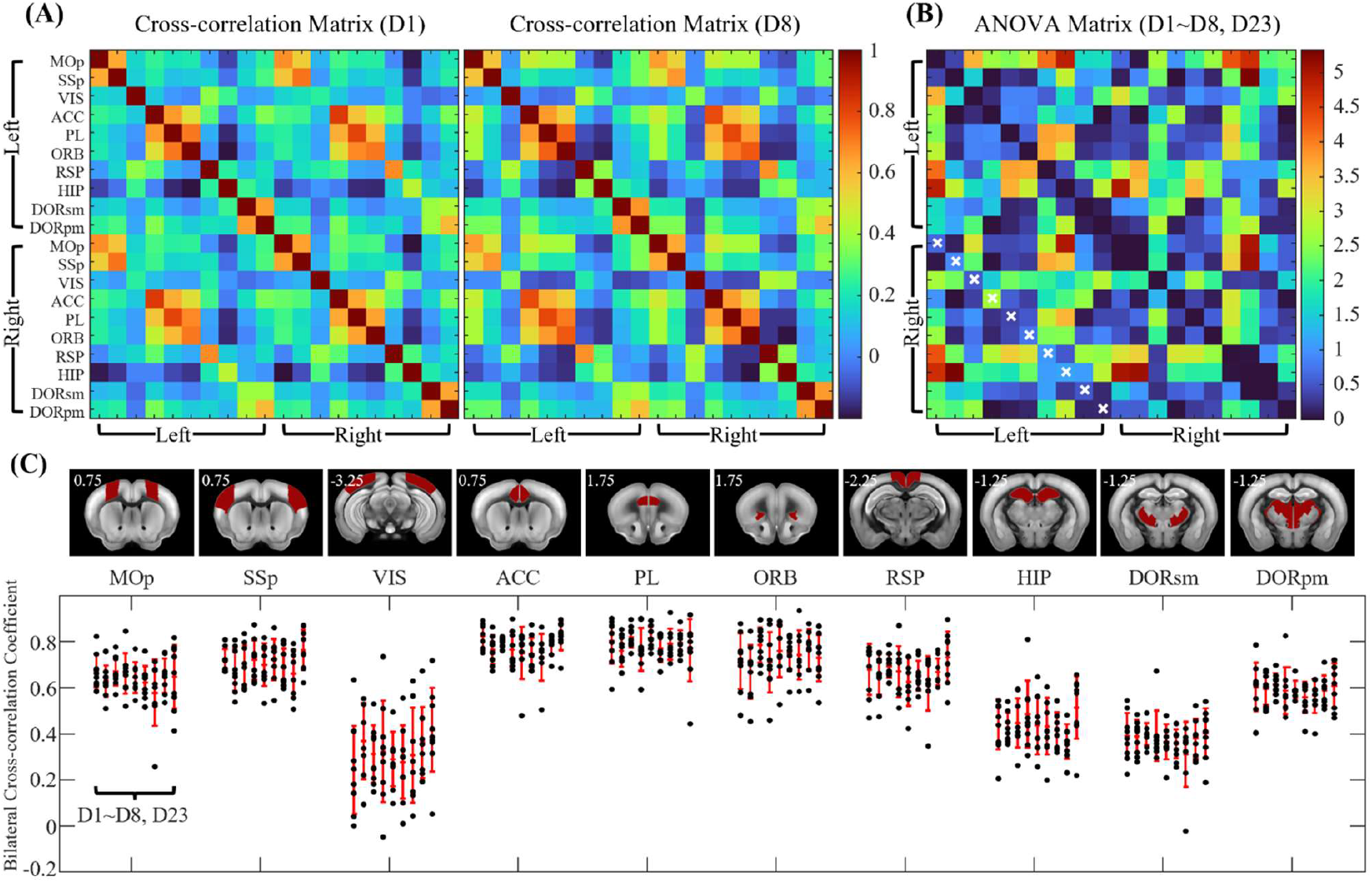
Cross-correlation matrices of 20 ROIs across the habituation period. **(A)** Signal correlation matrix of day 1 and day 8 datasets. **(B)** -log (p-value) matrix for ANOVA comparing day 1-8 and day 23, with p-values shown without FDR correction. No significant differences were detected after FRD correction. Bilateral region pairs are marked with ‘x’ in the matrix. **(C)** Bilateral cross-correlation changes across days 1-8, and day 23 for the ROIs (‘x’ marked region in c). No significant changes were detected in the ANOVA test. Ten regions of interest in each hemisphere are as follows: MOp, primary motor; SSp, primary somatosensory; VIS, visual; ACC, anterior cingulate; PL, prelimbic; ORB, orbital; RSP, retrosplenial; HIP, hippocampal; DORsm, thalamus sensory-motor cortex related; DORpm, thalamus polymodal association cortex related.

## 4 Discussion

Awake mouse fMRI is a promising tool for neuroscience and translational research. To minimize head motion during fMRI scanning, physical restraint is typically applied(see a review article, Mandino et al., 2024)[23], which can in turn induce stress. In our study, we found that stress levels slightly decreased with habituation outside the magnet but remained elevated during 8 days of habituation within the MRI environment. This suggests that mechanical vibrations associated with fast imaging may hinder habituation during the examined period. Head motion and functional connectivity remained stable across repeated fMRI sessions over the week, likely due to the well-designed restraint cradle that allowed paw movement.

### 4.1 Stress level

Multiple stressors are associated with awake mouse fMRI, including head and body restraint to minimize motion artifacts, as well as acoustic and vibrational noise. To reduce stress under these conditions, two primary strategies have been explored: (1) using less restrictive restraint systems, and (2) performing habituation under conditions that closely mimic the actual experimental environment.

#### Restraint

Regarding restraint, various head-mounting approaches have been developed, as summarized by Madino et al. (2024)[23]. Surgically implanted head-posts clearly provide more effective head stabilization than non-invasive restraint systems[23, 41]; therefore, we adopted a head-post implantation model. Although tilting the head at a 30° angle has been reported to reduce stress levels[25], we did not apply this configuration because the altered B_1_ field orientation relative to B_0_ can reduce MRI sensitivity and only a 3-cm radius space is available for our setup at 15.2 T.

In addition to head restraint, body restraint is often applied to further minimize motion-induced artifacts. In our design[16], we developed a cradle that provides a snug fit while allowing free paw movement, preventing the mouse from pushing against the restraint. Allowing paw movement has been shown to reduce both stress levels [25] and head motion[16].

#### Acclimatationd

Various habituation strategies have been adopted across laboratories [23]. In most cases, acoustic EPI noise was presented to mice during the habituation process; however, it did not appear to induce stress in C57BL/6 mice[25, 30], likely due to a mismatch between the dominant MRI acoustic frequency (peaking around ∼5 kHz) and the mouse hearing sensitivity range (peaking around ∼20 kHz)[25], as well as age-related hearing loss that occurs after approximately 8 weeks of age[42, 43]. Therefore, in our habituation protocol, acoustic noise was not applied, allowing acclimation without the need for a dedicated soundproof mock scanner room.

The daily duration and total number of habituation sessions are critical factors that vary widely across laboratories and experimental designs. In general, longer habituation sessions are often employed to reduce stress[23, 44]. However, repeated restraint can induce long-lasting physiological and neural alterations associated with stress, including changes in pain sensitivity[44] and modifications in hypothalamic gene expression related to stress regulation[45]. Therefore, we did not extend the habituation period beyond one week.

Regardless of the specific habituation strategy, assessing stress levels is essential-either indirectly through respiratory rate, heart rate, body weight, and food intake [17, 44, 46-49], or directly via plasma CORT measurements[17, 18, 25, 30, 41, 47]. Restraint stress typically reduces body weight and food intake while increasing respiratory rate, heart rate, and plasma CORT levels.

In our measurements, body weight decreased during the initial days of habituation, and plasma CORT levels increased to approximately 2-4 times the baseline level, indicating elevated stress.

CORT levels slightly decreased during one week of habituation outside the magnet, consistent with previous observations[25], but remained elevated during eight days of habituation inside the MRI scanner. Even after two weeks of habituation in a mock scanner, CORT levels increased to 2-3 times the habituated values during actual fMRI scanning (see Supplementary Fig. 6 in Xu et al., 2022; ∼150 ng/ml to ∼450 ng/ml)[25], which is comparable to our observations without prior habituation (216 ng/ml to 628 ng/ml). These findings suggest that mechanical vibration is a major stressor that cannot be reproduced outside the magnet and is not easily habituated. This further implies that even when mice are well-trained to perform cognitive tasks in a mock scanner, their task performance may be reduced during actual fMRI scanning due to the additional vibration-induced stress.

To reduce mechanical vibrations and acoustic noise associated with EPI scanning, one potential approach is to employ acoustically silent zero-echo-time (ZTE) or ultrashort-echo-time (UTE)-based pulse sequences[50]. However, since their fMRI contrast partly arises from inflow effects induced by rapidly repeated low-flip-angle RF pulses[51], their application is generally restricted to setups using a surface coil for both transmission and reception, rather than the more commonly used body excitation coil.

### 4.2 Impact of habituation to rsfMRI

Head motion is an important indicator of both stress and fMRI data quality. In our experiments, head motion remained similar across all sessions, with FD values of approximately 29 μm, regardless of habituation. This value is slightly higher than that reported in our previous study (20 μm FD)[16]. The exact source of this difference is unclear but may reflect variability across animal batches.

Although acclimation is generally expected to reduce head motion[18], we did not observe decreases in head motion over days, consistent with the lack of changes in stress-related measures. This is consistent with recent observations comparing 4-day and 13-day habituated mice (Fig. 2c in Laxer et al, 2025)[41].

To reduce motion-related artifacts in fMRI, data scrubbing is often applied in awake fMRI analyses. However, because our goal was to compare time-dependent rsfMRI functional connectivity across days, no data scrubbing was applied, nor sophisticated data processing procedures were adopted. In our dataset, both FC specificity and FC between ROIs remained stable before, during, and after the habituation period. This indicates that the observed stability in functional connectivity reflects consistent neural network across the habituation process.

## 6 Author Contributions

SH Choi, GH Im and SG Kim conceived the ideas and designed the experiments, GH Im performed all experiments, SH Choi processed all imaging-related data, and SH Choi and SG Kim wrote the manuscript.

## 8 Acknowledgments

We thank Junglim Lee for providing animal care and performing CORT measurements.

