## Supplementary Figure 1 for "Longitudinal Resting-State fMRI of Awake Mice During Habituation: Stress, Head Motion, and Functional Connectivity": Supplementary_Material.pdf

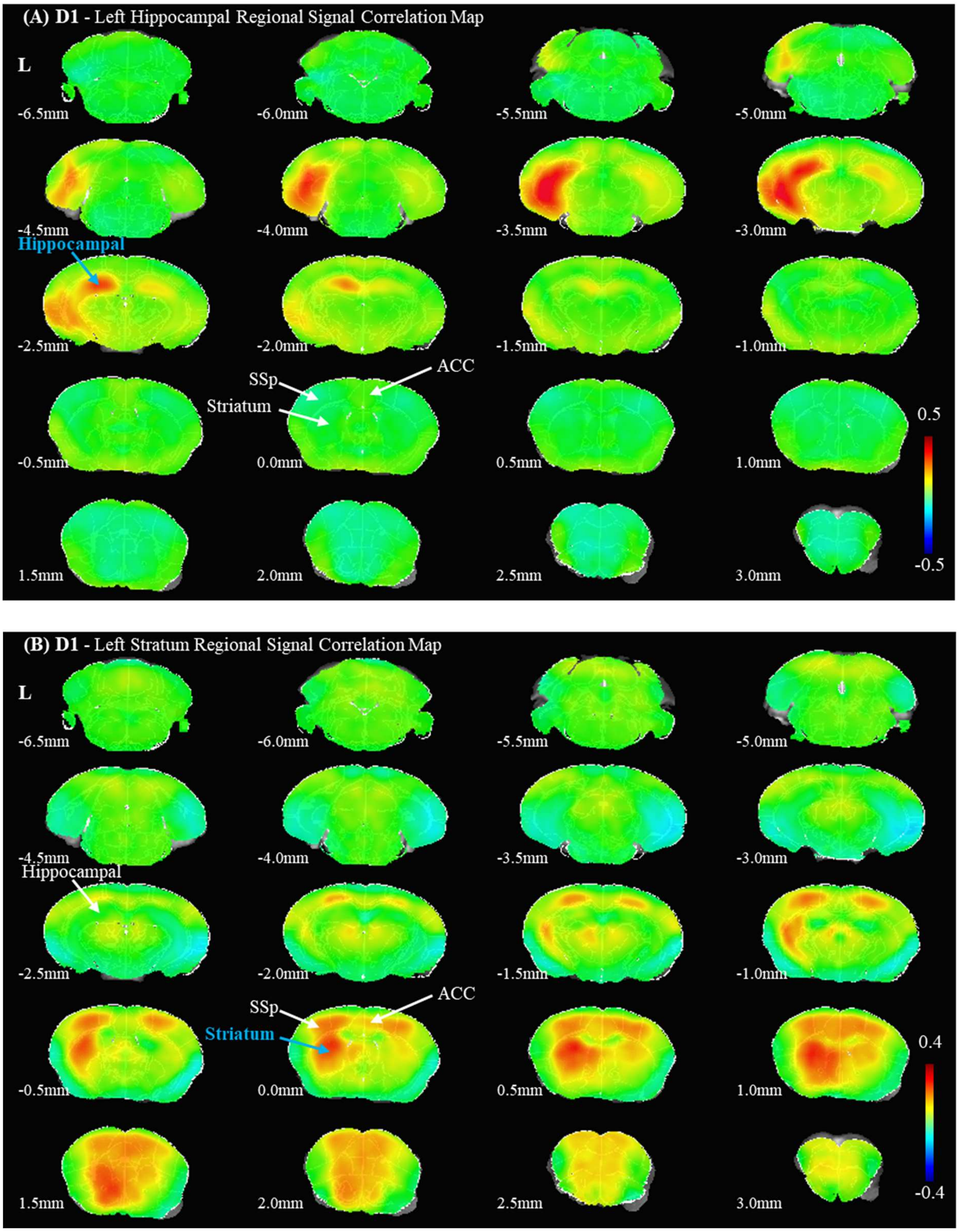


**Supplementary Figure 1. Hippocampal region- and Striatum-seed based cross-correlation maps of the resting-state fMRI on the initial habituation day (n = 9).** (A) Cross-correlation maps based on the left hippocampal seed. (B) Cross-correlation maps based on the left striatal seed. ACC, anterior cingulate; SSp, primary somatosensory.
